# Early categorization of social affordances during the visual encoding of bodily stimuli.

**DOI:** 10.1101/2022.09.29.510147

**Authors:** Q. Moreau, E. Parrotta, U.G. Pesci, V. Era, M Candidi

## Abstract

Interpersonal interactions rely on various communication channels, both verbal and non-verbal, through which information regarding one’s intentions and emotions are perceived. Here, we investigated the neural correlates underlying the visual processing of hand postures conveying social affordances (i.e., hand-shaking), compared to control stimuli such as hands performing non-social actions (i.e., grasping) or showing no movement at all. Combining univariate and multivariate analysis on electroencephalography (EEG) data, our results indicate that occipito-temporal electrodes show early differential processing of stimuli conveying social information compared to non-social ones. First, the amplitude of the Early Posterior Negativity (EPN, an Event-Related Potential related to the perception of body parts) is modulated differently during the perception of social and non-social content carried by hands. Moreover, our multivariate classification analysis (MultiVariate Pattern Analysis - MVPA) expanded the univariate results by revealing early (<200ms) categorization of social affordances over occipito-parietal sites. In conclusion, we provide new evidence suggesting that the encoding of socially relevant hand gestures is categorized in the early stages of visual processing.

## Introduction

Humans spend much of their lives observing how their conspecifics act, predicting and understanding their intentions, and learning from them. These reciprocal processes are crucial to efficient inter-individuals relationships. Specifically, during interpersonal interactions, non-verbal information conveyed by facial expressions, body postures, and hand movements represents a key channel through which others express their intentions and their willingness to make connections (Argyle, 1972).

Based on evidence from neurophysiological (Thierry *et al*., 2006; Meeren et al., 2013), neuroimaging (fMRI, Downing *et al*., 2001; Gschwind *et al*., 2012), brain lesion (Sliwinska *et al*., 2020), and brain stimulation studies (Urgesi *et al*., 2007; Sadeh *et al*., 2011), a specific visual pathway for the processing of social stimuli such as faces, bodies, and their movements, has recently been proposed (Pitcher and Ungerleider, 2021). Located on the lateral surface of the occipital and parietal lobes, this third-visual “social “ pathway is an addition to the two-streams model (i.e., the “what “ ventral pathway and the “where “ dorsal pathway, Milner and Goodale, 2006; Milner and Goodale, 2008; Goodale, 2014).

While most seminal neurocognitive models have focused on face perception and its contribution to sociality (see Haxby *et al*., 2000; Haxby *et al*., 2002; Hugenberg and Wilson, 2013 for reviews), body postures and hand gestures also constitute a highly relevant source of social information for efficient interpersonal interactions (Ramsey, 2018). The processing of body images has been associated with the activity of a bilateral occipito-temporal focal region, the Extrastriate Body Area (EBA; Downing *et al*., 2001), whose causal role in visual body perception has been confirmed by brain stimulation and lesion studies (Urgesi et al., 2004; Urgesi *et al*., 2007; Moro *et al*., 2008; Moro *et al*., 2012; Gandolfo and Downing, 2019). Interestingly, imaging studies showed a topographical functional organization of high-order temporo-occipital visual areas (Orlov *et al*., 2010), with specific brain regions of this territory associated with the processing of hand images (compared to other body parts) in the lateral occipital-temporal cortex and the extrastriate body area (i.e., LOTC and EBA; Bracci *et al*., 2010; Bracci *et al*., 2012). Patterns of activity of the LOTC are thought to form a representational space encompassing low- (e.g., body-related shapes) and high-level (action type, social meaning) aspects of social vision (Lingnau and Downing, 2015; Wurm *et al*., 2017; Tucciarelli *et al*., 2019; Tarhan and Konkle, 2020). Non-invasive brain stimulation and lesion studies showed that the discrimination and recognition of implied (social and non-social) actions further depend on the activity of superior temporal (Candidi *et al*., 2011; Urgesi *et al*., 2014; Candidi *et al*., 2015), parietal (Cattaneo *et al*., 2010; Avenanti *et al*., 2013), and premotor (Urgesi *et al*., 2007; Candidi *et al*., 2008; Cattaneo *et al*., 2010; Avenanti *et al*., 2013; Urgesi *et al*., 2014) regions.

The early role of occipito-temporal areas in action perception is currently debated. Especially the pending interrogation remains to what extent the LOTC alone can handle the encoding and the discrimination of actions, and whether and how this activity is later supported and integrated within the fronto-parietal network (Lingnau & Downing, 2015; Tucciarelli *et al*., 2019; Tarhan *et al*., 2020). Indeed, to the best of our knowledge, only one MEG study (Tucciarelli *et al*., 2015) investigated the spatio-temporal dynamics of action discrimination and indicated that the activity of lateral occipito-temporal regions (i.e., consistent with the EBA) has the earliest access (compared to precentral regions) to the discrimination of different observed actions (e.g., grasping and pointing) independently from their low-level features. Furthermore, previous fMRI studies (Wurm *et al*., 2017; Bracci *et al*., 2018) have investigated how the LOTC processes hand stimuli in a view-invariant manner, since this is a computationally fundamental characteristics in order to recognize different postures conveying specific information, as well as along features of sociality. Yet, no study has tested the temporal dynamics of *social* action representation, crucial to the understanding of how social affordances are encoded by the visual cortex.

The neurophysiological response of occipito-temporal areas to the presentation of body (e.g., hand) images has been characterized in the time domain by Magneto-Electro-, and intracortical-physiological recording studies (i.e., M/EEG/ECoG) showing that the visual perception of hands modulates the amplitude of a negative component (i.e., the N190) over occipito-temporal regions (Thierry *et al*., 2006; Ishizu *et al*., 2010). Interestingly, the perception of hand images elicits a higher amplitude of the N190 compared to both whole bodies, other body parts and control images (Espírito Santo *et al*., 2017; Moreau et al., 2018; Moreau *et al*., 2019). Studies also showed that the socio-emotional content of bodily stimuli could modulate the amplitude of the N190. For instance, Borhani and colleagues (2015) report a larger N190 component when images of fearful bodies are perceived, compared to other emotions and control conditions. On the other hand, other studies have reported no N190 amplitude modulation during the perception of emotional (e.g., insulting) gestures (Flaisch and Schupp, 2013). Altogether, it is important to note that the neural mechanisms underlying the N190 amplitude modulation in response to emotional bodily stimuli are still a topic of ongoing research. Furthermore, only a few studies have directly studied the effect of socially meaningful body postures compared to other types of movements on early visual responses (Flaisch *et al*., 2011, 2013) and a parallelism and/or distinction between emotionally and socially meaningful postures is not straightforward.

Besides these early and posterior brain responses to visual bodily stimuli, EEG studies have also highlighted the sensitivity of later and more cognitive brain responses such as the Early Posterior Negativity (EPN), peaking around 300 ms after stimulus presentation, to salient stimuli. So far, EPN has been studied during the visual processing of socially relevant stimuli such as emotional faces (Sato *et al*., 2001) and body postures (Borhani *et al*.,2015), and its modulation is thought to reflect attentional selection to relevant stimuli that will generate further processing (Schupp, Junghöfer, *et al*., 2004).

In the present study, we aimed to describe the neural dynamics associated with perceiving hand stimuli that convey social affordances. We recorded EEG activity during the passive viewing of hand pictures either being still, performing a non-social action (i.e., grasping), or conveying a social affordance (i.e., hand-shaking; Figure 1A). Our main hypothesis was that the visual perception of hands conveying social affordances would generate different EEG signatures (i.e., N190 and EPN) relative to hands not providing any social affordance. To do so, classical univariate (i.e., event-related potentials - N190 and EPN) and multivariate (Multivariate Pattern Analysis - MVPA) analyses were combined to explore the processing of socially relevant hand postures. This approach has recently gained momentum to get new insights into the neural patterns of activity underlying perceptual and cognitive processes across multiple scales (Grootswagers *et al*., 2017).

**Figure 1.**
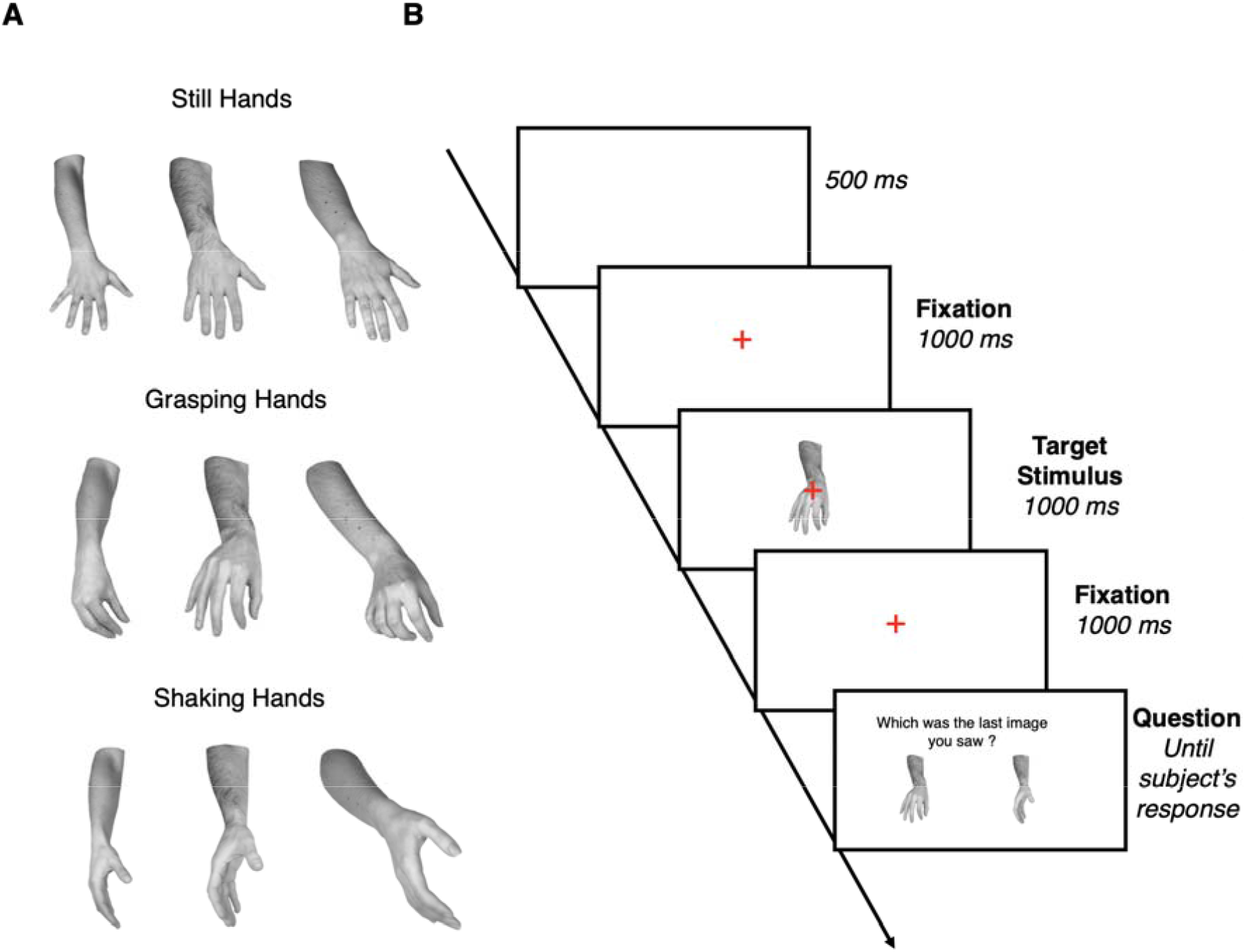
**A**) Set of stimuli for each of the three classes. **B**) Experimental timeline.

## Material and Methods

### Participants

23 healthy individuals (10 males and 13 females, mean age 25.3 years) took part in the experiment. The sample size was established using a power analysis, performed with the software More Power (Campbell and Thompson, 2012). We inserted as expected effect size the partial eta squared value obtained by Moreau *et al*. 2019 (η_p_^2^ = 0.29). The output indicates that a three-level factor in a within-subject design, with a power of 0.95 and a partial eta squared of 0.29, requires a sample size of 22 participants.

All participants were right-handed with normal or corrected-to-normal vision acuity in both eyes. Information about the purpose of the study was provided only after the experimental tests were completed. Participants gave their written informed consent and were reimbursed at a rate of 7€/hour. The experimental procedures were approved by the Ethical Committee of the Fondazione Santa Lucia (Rome, Italy) and were in accordance with the ethical standards of the 2007 Declaration of Helsinki.

### Stimuli

Original stimuli comprised a set of 12 grayscale pictures taken with a digital camera in the same space and light intensity-controlled conditions for each class of stimuli. The stimuli set was made of three types of hand pictures: 1) Still hands (no action), 2) Grasping hands conveying an implied action without social affordance, and 3) Shaking hands conveying an implied action with social affordances (Figure 1A). Each category comprised a set of four images from four different volunteers ‘ right hands (2 males, 2 females). The images were modified by means of the Adobe Photoshop software to obtain a set matched for dimension, orientation, background, contrast, and luminosity; all images were scaled to 400×400 pixels at a resolution of 72 dpi on ‘Netscape grey ‘ (128;128;128).

Stimuli have been validated by two online surveys beforehand (Survey Monkey) to control for both clarity (distinguishing the specific movement) and socially informative features (distinguishing the presence/absence of social cues), as well as rates of implied motion conveyed by the stimuli (Grasping and Shaking hands). In the first survey, 30 individuals were required to evaluate the original set of 22 images, by assessing the degree of implied motion conveyed by each image (Likert scale from 1 to 10), as well as the meaning of the implemented movement (open question: “describe, in your own words, what is the hand doing). Overall, 61.49% of the final stimuli set was accurately described (i.e., open description mentioning grasping, shaking, or absence of movement in the hands ‘ pictures), and the rates of implied motion assessed by a t-test revealed no difference between Grasping and Shaking images (*t*(29) = 1.625, *p* = 0.12) while the implied motion was lower in Still hands than both Grasping hands (*t*(29) = 9.373, *p* < 0.001) and Shaking hands (*t*(29) = 11.144, *p* < 0.001). In the second survey, we aimed to further validate the selected stimuli. We asked another 30 participants to answer two questions for each stimulus. First, they had to complete the sentence “the action performed by the hand is directed towards: … “ and choose between “an object “, “another person “, or “neither of the two “. Grasping hands were correctly identified as moving towards an object in 90.83% of the cases, shaking hands were correctly identified as moving towards another person in 90.00% of the cases, while Still hands seemed to be perceived as more ambiguous, with 53.33% of the participants correctly answering “neither of the two “. Second, we explicitly asked whether the hand displayed was still, grasping an object or shaking another hand. Participants correctly answered that our Grasping hands ‘ stimuli were grasping an object (95%), that our Social hands were shaking another hand (84.16%), and that our Still hands were motionless (83.33%).

An analysis of the ERPs for each hand category but separately for each of the four sets showed that images from one of the four volunteer subsets generated a very different pattern of results in the Grasping condition compared to the other three (See Supplementary Figure S1). To avoid any confounding effect in the interpretation of the data, and in order to keep the numerosity of the different conditions balanced, we have decided to remove trials associated with the presentation of the subset of images of this identity. Therefore, the final stimuli set comprised 9 grayscale pictures (3 Still, 3 Grasping, and 3 Shaking hands).

### Procedure

Each trial consisted of the presentation of a red fixation cross for 1000 ms before the appearance of the target stimulus. Target images were shown for 500 ms at the center of the screen. The fixation cross remained on the screen during the presentation of the target stimulus in order to avoid EEG artifacts associated with its appearance and disappearance. Stimuli were viewed at a distance of approximately 60 cm. In order to maintain subjects ‘ attention on the stimuli, after the disappearance of the stimulus, in 30% of the trials, participants were asked to choose one among two presented images by pressing one of the two keys from the pad with their right hand to indicate which stimulus they thought was the one presented in that trial (Figure 1B). Stimuli presentation was displayed by using E-Prime 2.0 and was randomized across subjects. The experiment consisted of three blocks with 100 trials each, where stimuli were presented randomly. The total number of trials was 300 (100 trials for each of the three different conditions).

### EEG-recordings and preprocessing

EEG signals were recorded and amplified using a Neuroscan SynAmps RT amplifiers system (Compumedics Limited, Melbourne, Australia) and acquired from 56 tin scalp electrodes embedded in a fabric cap (Electro-Cap International, Eaton, OH), arranged according to the 10-10 system. The EEG was recorded from the following channels: Fp1, Fpz, Fp2, AF3, AF4, F7, F5, F3, F1, Fz, F2, F4, F6, F8, FC5, FC3, FC1, FCz, FC2, FC4, FC6, T7, C5, C3, C1, Cz, C2, C4, C6, T8, TP7, CP5, CP3, CP1, CPz, CP2, CP4, CP6, TP8, P7, P5, P3, P1, Pz, P2, P4, P6, P8, PO7, PO3, POz, PO4, PO8, O1, Oz, O2. Horizontal electro-oculogram (HEOG) was recorded bipolarly from electrodes placed on the outer canthi of each eye and signals from the left earlobe were also recorded. All electrodes were physically referenced to an electrode placed on the right earlobe and were algebraically re-referenced offline to the average of both earlobe electrodes. Impedance was kept below 5 KΩ for all electrodes for the whole duration of the experiment, the amplifier hardware band-pass filter was 0.01 to 200 Hz and the sampling rate was 1000 Hz. In order to remove the eye blinks or other artifacts, a blind source separation method was applied (i.e. Independent Component Analysis - ICA; Jung *et al*., 2000) implemented using the FieldTrip routines (Donders Institute, Nijmegen, The Netherlands; Oostenveld *et al*., 2011) to remove from the EEG any components related to eye movements. Then, EEG data were band-pass filtered (high pass = 0.5, low-pass = 40 Hz) and segmented into epochs of 8000 ms from -4000 ms to +4000 ms around stimulus onset. Finally, all trials were visually inspected in order to further remove artifacts related to amplifier blocking, residual eye blink, or muscular noise.

### ERP analysis

Analyses of P1, N190, and EPN components were performed in MATLAB R2019a using the FieldTrip routines (Donders Institute, Nijmegen, The Netherlands; Oostenveld *et al*., 2011). Before ERP averaging across subjects for each condition, EEG epochs were baseline corrected from -200 ms to 0 ms before stimulus presentation. Based on previous studies (Borhani *et al*., 2015, Thierry *et al*., 2006, Schupp *et al*., 2006, Moreau *et al*., 2018, Moreau *et al*., 2020) and the visual inspection of the data, the P1 amplitude was quantified as the mean amplitude in a time window of 110-170 ms post-stimulus onset, the N190 amplitude as the mean amplitude in a time window of 160-230 ms post-stimulus onset, and the EPN as the mean amplitude in a time window of 250-350 ms post-stimulus onset. Each component ‘s mean amplitudes were further analyzed through a one-way within-subject repeated measures ANOVA with CATEGORY as a factor (i.e., Still, Grasping and Shaking hands) in R (R Core Team, 2014) using the ez-package (Lawrence, 2016). Mean amplitude values were extracted over bilateral occipito-temporal clusters of electrodes (i.e., P7/8 and PO7/8, see (Borhani *et al*., 2015; Moreau *et al*., 2018; Moreau *et al*., 2019). The Greenhouse-Geisser correction for non-sphericity was applied when appropriate (Keselman *et al*., 1980), and post-hoc comparisons were performed using the Bonferroni correction for multiple comparisons.

### Multivariate Pattern Classification Analysis

#### Preprocessing

The EEG data used for the classification were low-pass filtered at 30 Hz (van Driel *et al*., 2021), segmented into epochs of 700 ms (from -200 ms to +500 ms around stimulus onset) in order to reduce computation time, and downsampled to 500 Hz. Further preprocessing steps as well as all the classification analyses were performed using the MVPA-Light MATLAB toolbox (Treder, 2020). Prior to classification, data were normalized using z-scores to center and scale the training data, providing numerical stability (Treder, 2020). Across all classification analyses, if a class had fewer trials than another, we corrected the imbalance by undersampling the over-represented condition (i.e., randomly removing trials). Then, using the preprocessing option *mv_preprocess_average_samples*, training data (i.e., EEG trials) from the same class (i.e., Hand Category) were randomly split into 4 groups and averaged, so that the single-trial dataset was replaced by 4 averaged trials (i.e., samples). The classifications were then run using these averaged samples, as this step is known to increase the signal-to-noise ratio (Grootswagers *et al*., 2017; Smith and Smith, 2019; Treder, 2020).

#### Classification in time

We performed 3 separate binary classification analyses in time (−200 to 500 ms around stimulus onset), using Still/Grasping, Still/Shaking, and Grasping/Shaking samples. To do so, a Linear Discriminant Analysis (LDA) classifier was implemented using the MVPA-Light toolbox. For linearly separable data, an LDA classifier divides the data space into *n* regions, depending on the number of classes, and finds the optimal separating boundary between them using a discriminant function to find whether data fall on the decision boundary (i.e., 50% classification accuracy) or far from it (i.e., > 50% classification accuracy). We used 10-fold cross-validation repeated 10 times on 4 averaged samples per class in order to quantify our classifier ‘s performance in terms of accuracy. At the group level, statistical significance (i.e., performance significantly above 0.5 chance level) was assessed through non-parametric cluster-based permutation tests (Maris and Oostenveld, 2007), using ‘maxsum ‘ for cluster corrections and Wilcoxon test for significance assessment. At the individual level, we also computed peak decoding, defined as the maximum accuracy value between 100 and 400ms after stimulus presentation, and peak decoding latency, corresponding to the time-point to which the maximum peak was detected (Mares *et al*., 2020). Both peak decoding and peak decoding latency variables were analyzed through a one-way ANOVA with CATEGORY as a factor in R using the ez-package.

#### Temporal generalization

In order to further understand how the different EEG patterns related to each binary classification evolve over time, we adopted the temporal generalization method (King & Dehaene, 2014). Specifically, we tested whether the neural representations carrying the classifier performance in each classification were stable (generalizable) across time, or if they rather vary over time. This approach allows testing whether the neural code that supports above-chance decoding is stable or is dynamically evolving. Instead of applying a different classifier at each time point, the classifier trained at time t can be tested on its ability to generalize to time t ‘. Generalization implies that the neural code that is identified at time t recurs at time t ‘. In order to do so, for each binary decoding, we trained our classifier on each single time point and tested it at all time points in the same time window previously selected for the classification across time (i.e., from -200 to 500 ms around stimulus onset). This provided a temporal generalization matrix (training time x generalization time) with accuracy values tested for statistical significance (Figure 4, lower panel). As proposed by King & Dehaene (2014), if a classifier generalized from one time-window to another, then the underlying processes for the two time-windows would be similar, and the values in the resulting training time x generalization time plot would be represented as a single area spreading continuously out of the diagonal. If this were not the case, the results would be based on different information that travels across a chain of transient neural representations, and the results in the training time x generalization time matrix would yield a diagonal generalization pattern.

#### Classification across time and electrodes

Using identical settings for the classification in time, we performed a spatio-temporal search using two neighboring matrices for time points and electrodes, respectively. Thus, in this analysis each electrode/time-point and its direct neighbors acted as features for the classification, resulting in a channel x time point matrix of accuracy scores. By plotting this matrix, we then assessed which electrode carried the highest weight in the temporal decoding (see Figure 5). Furthermore, we used Fieldtrip-implemented cluster-based analysis (Maris and Oostenveld, 2007) to find significant differences in the topographical distribution of classification weights over time between the three binary decodings we have run.

## Results

### Behavioral results

Overall, subjects responded correctly at 99.326 % when asked to identify the stimulus presented.

### Event-Related Potentials

The results of the ERP analysis are shown in Figure 2.

**Figure 2.**
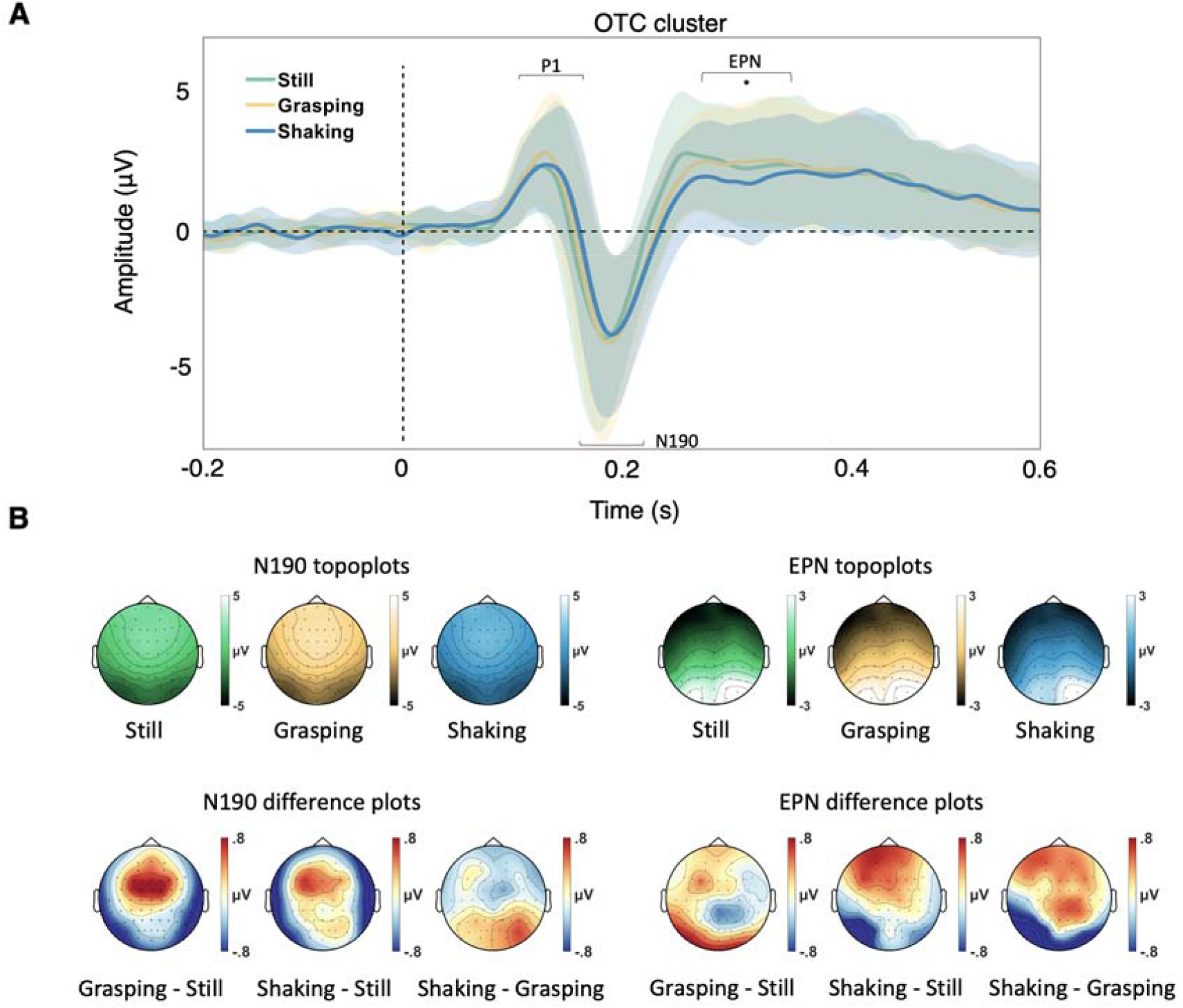
**A**) P1, N190 and Early Posterior Negativity (EPN) event-related potentials over the bilateral occipito-temporal cluster (OTC; P7, PO7, P8, and PO8) for the three conditions, shaded lines represent standard deviation; **B**) Upper panels: topographical plots of the N190 (160-230 ms) and the EPN (250-350 ms) for the three conditions; lower panels: topographical plots of the difference waves between conditions (i.e. Grasping minus Still, Shaking minus Still, Shaking minus Grasping) in the N190 and EPN time windows.

*P1 component*. The ANOVA on the P1 mean amplitudes revealed no significant difference between conditions [*F*(2,44) = 1.41, *p* = 0.25, η_p_^2^ = .060].

*N190 component*. The ANOVA on the N190 mean amplitudes revealed no significant difference between conditions [*F*(2,44) = 1.61, *p* = 0.21, η_p_^2^ = .068].

*EPN component*. The ANOVA on the EPN mean amplitudes revealed a main effect of CATEGORY [*F*(2,44) = 8.49, *p* <.001, η_p_^2^ = .278] over the bilateral occipito-temporal cluster (P7, PO7, P8, and PO8). Bonferroni corrected post-hoc comparisons revealed a significantly larger EPN amplitude associated with Shaking hands (M = 1.8µV, SD = 1.98) compared to Grasping hands (M = 2.4µV, SD = 1.87) [*p* = .008] and Still hands (M = 2.54µV, SD = 2.06) [*p* = .001, see Figure 2] and no significant difference between Still and Grasping hands [*p* = 1].

### Classification in time

To test whether implied motion and social information displayed in hand stimuli were differently reflected in the EEG time series, we trained and tested a classifier on the EEG data associated with the presentation of those images. Our linear classifier was trained to discriminate between A) Still and Grasping hands, B) Still and Shaking hands, and C) Grasping and Shaking hands. The results from this analysis, plotted as classification accuracy (ACC), are shown on the upper panels of Figure 3. Statistically significant results (*p* < .05, classification chance level 0.5) are highlighted by a bold line. First, all classifications show at least one significant cluster, meaning that our classifier was able to discriminate between conditions above chance level. The Still vs Grasping classification shows the lowest performance values and the classifier reaches significance almost 100 ms later than in the Still vs Shaking classification (i.e., from 220 ms compared to 132 ms, significant clusters time window: [220-264 ms; 280-308 ms; 346-386 ms; 448-476 ms]). The highest accuracy values are observed for the Still vs Shaking classification (i.e., 0.61 at 152ms, significant clusters time window: [132-328 ms; 340-406 ms; 424-500 ms]). Interestingly, in the Grasping vs Shaking decoding, the classifier ‘s performance reaches significance later than in the other classifications (significant clusters time window: [282-340 ms]). Furthermore, the one-way ANOVA on peak decoding revealed a main effect of CLASSIFICATION (*F*(2,44) = 3.451, *p* = 0.041, η_p_^2^ = .136). Post-hoc analysis using the Bonferroni correction for multiple comparisons showed that peak decoding values were higher in the Still vs Shaking decoding than in Still vs Grasping one (*p* = 0.036 - Figure 3C). The ANOVA on peak decoding latency revealed no significant difference between classifications (*F*(2,44) = 1.372, *p* = 0.264, η_p_^2^ = .059 - Figure 3D).

**Figure 3.**
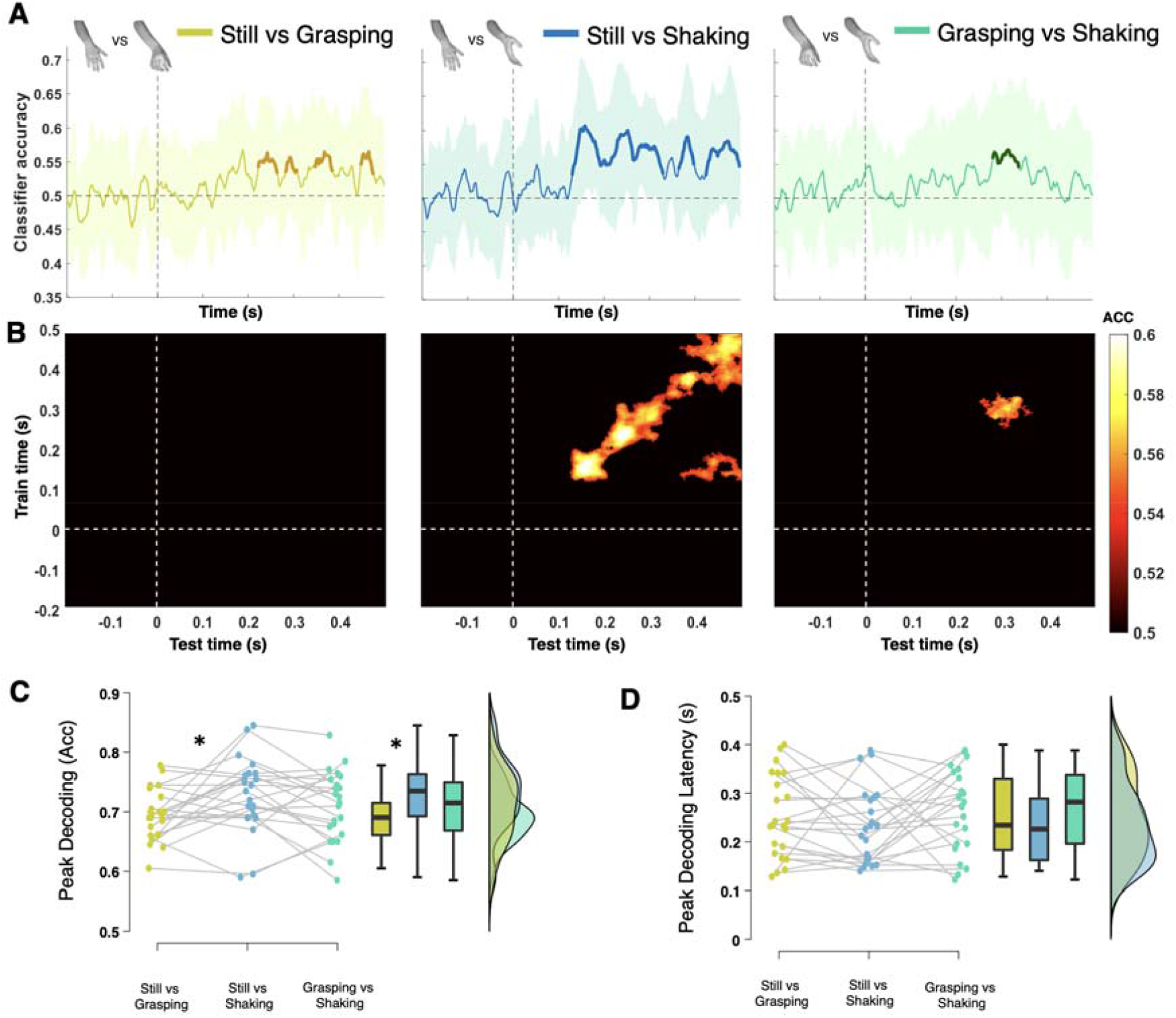
**A)** Temporal decoding and significant clusters for the binary comparison of Still vs Grasping, Still vs Shaking and Grasping vs Shaking. Shaded lines represent standard deviation. **B)** Temporal generalization matrix for each binary comparison, masked to show only significant (*p* > .05) accuracy scores. **C)** Peak decoding and **D)** peak decoding latency raincloud plots across the three binary decodings.

### Temporal generalization

Since our decoding analysis across time (Figure 3A, upper panels) showed that the classifier ‘s performance reached significance at different timings for each binary comparison and that the periods of significant decoding were more sustained during Still vs Shaking than during Still vs Grasping and Grasping vs Shaking, we run a temporal generalization analysis. This approach allows testing whether the neural code that supports above-chance decoding is stable or is dynamically evolving. Instead of applying a different classifier at each time point, the classifier trained at time t can be tested on its ability to generalize to time t ‘. The resulting training time x generalization time matrices (Figure 3B, masked to show only significant (p < .05) results) show whether the classifier relied on the same or on different neural patterns across time when successfully classifying between classes. Overall, the classifier ‘s performance highlights that the classification in each binary comparison relied on different patterns. In particular, we observe that when the classifier had to distinguish between Still and Shaking hands its performance did not generalize over time (Figure 3B, central panel). Rather, the diagonal generalization pattern suggests that the processes leading to each time-window of classification (in particular the first until ∼200 ms and the second from ∼200 to 300 ms) are different. Interestingly, in the Still vs Shaking comparison, we find only one cluster outside the diagonal (from ∼350 to 500 ms). This might suggest that, when classifying Still vs Shaking hands, later generators reactivate the ones involved during the first time window of significant classification (King & Dehaene, 2014).

### Classification in time and space

We performed the same binary comparisons used for the above time classification analysis over time and electrodes (adopting a similar “saliency map “ approach, as reported by Vahid *et al*., 2020) (Figure 4 A-C) with the same parameters and statistical analysis (i.e., LDA classifier, cluster-based permutation test correcting for multiple comparisons). This additional ‘searchlight ‘ analysis highlights which EEG electrodes are most important for the classification (Treder, 2020). The results are shown in Figure 4 A-C, where only the significantly above chance level (*p* < .05, classification chance level 0.5) points are displayed. Overall, the time and space decoding are in line with the temporal results but reveal different spatial patterns. We observe a distributed activation pattern across the scalp (i.e., occipital, centroparietal, frontocentral, Figure 4B) when distinguishing between Still hands and Shaking hands, while the decoding between Still and Grasping, as well as the one between Grasping and Shaking, only seem to rely on parieto-occipital sites (Figure 4A and 4C).

**Figure 4.**
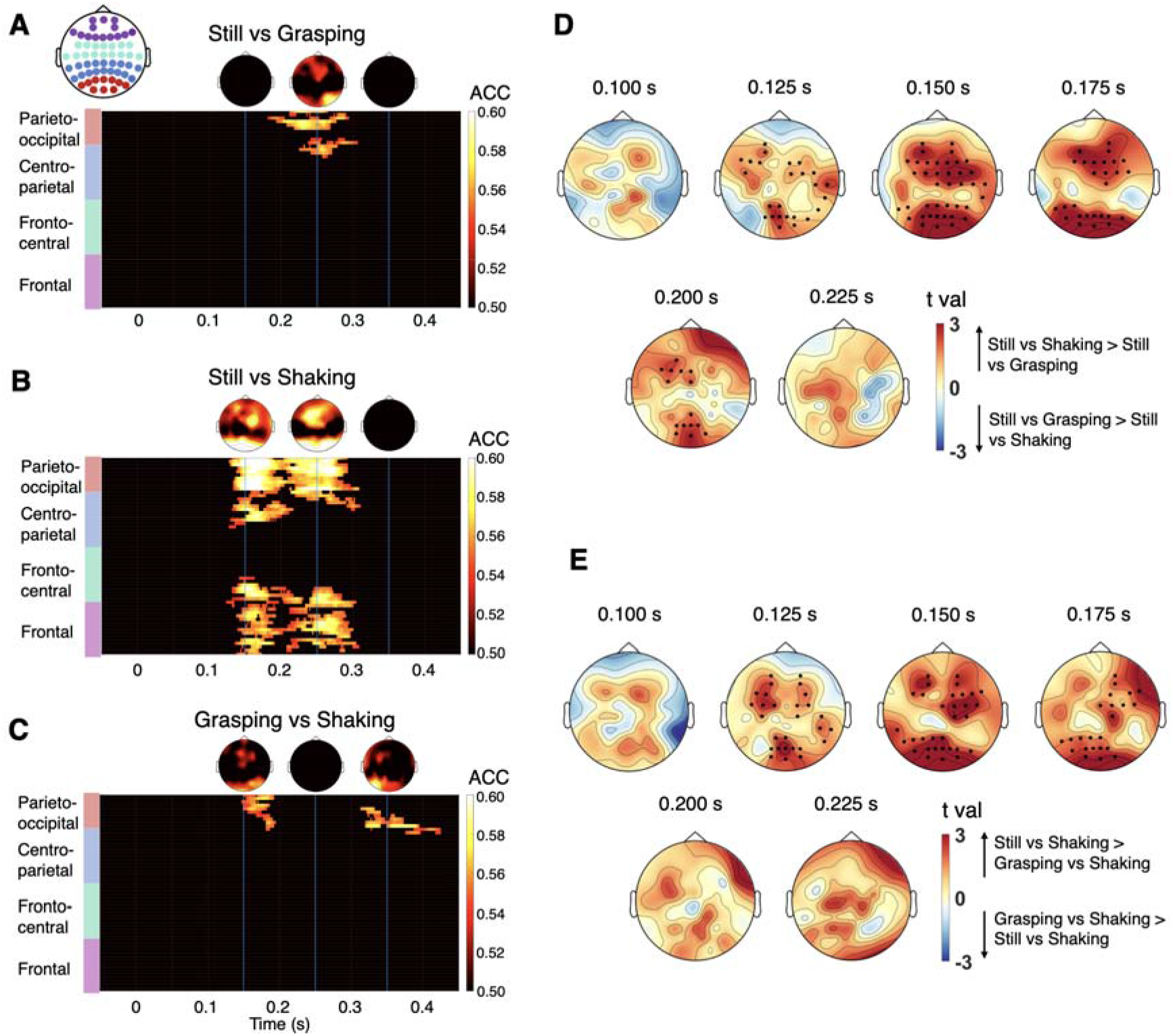
Spatio-temporal decoding and significant clusters for the binary comparison of **A**) Still vs Grasping; **B**) Still vs Shaking and, **C**) Grasping vs Shaking. Each row of the heatmap represents one EEG channel, the channels are grouped regarding their spatial distribution over the scalp (Parieto-Occipital, Centro-Parietal, Fronto-Central, Central). The topographies above each plot show ACC values at three points in time, highlighted by the three light blue bars (i.e. 150, 250 and 350 ms). Results of the cluster-based permutation test showing significant (cluster *p* < .05) differences in the topography of classification over time between **D)** the Still vs Shaking and Still vs Grasping decoding, and **E)** the Still vs Shaking and Grasping vs Shaking decoding. No significant differences were found between the Still vs Grasping and Grasping vs Shaking decoding.

### Cluster-based permutation test comparing the classification topographies for each time x electrode binary decoding

Finally, in order to find significant differences in the topographical distribution of classification weights over time between the three binary decodings we have run, we have conducted a cluster-based permutation test (Maris and Oostenveld, 2007). Coherently with what we observe in Figure 4 A-C, Figures 4D and 4E show that the Still vs Shaking decoding is characterized by different patterns compared to Still vs Grasping and Grasping vs Shaking. The cluster-based results emphasized this, showing that there were no clusters of significant differences (*p* > .05) when comparing the performance of Still vs Grasping and Grasping vs Shaking, whereas Still vs Shaking significantly differed from the other two, being characterized by an earlier classification onset (from 125 ms) as well as by a fronto-central cluster of electrodes contributing to the classification (*p* < 0.001 for Still vs Shaking VS Still vs Grasping; *p* < 0.005 for Still vs Shaking VS Grasping vs Shaking).

## Discussion

The aim of the present study was to investigate the EEG markers associated with the visual perception of hand gestures conveying social affordances (shaking hands). By using control stimuli showing hands performing non-social actions (i.e., grasping actions, matched for implied motion with shaking hands) and still hands, we combined univariate analyses of the P1 component, the early body-selective N190 (Thierry *et al*., 2006) and mid-latency attentional selection EPN (Borhani *et al*., 2015) indexes with multivariate analyses to give a detailed description of the neural (temporal and spatial) dynamics at play during the processing of hand images containing social information. Describing the spatio-temporal dynamics of social affordances visual processing represents a key step, contributing to the current investigation of the neural underpinnings of social vision (Lingnau & Downing, 2015; Tucciarelli *et al*., 2019; Tarhan *et al*., 2020, Papeo, 2020).

### ERPs modulations in response to hand gestures conveying social affordances

First, our univariate analysis reveals that neither the P1 nor the early body-selective N190 component is significantly influenced by the social meaning of the movement implied in hand postures. The N190 component has traditionally been associated with the encoding of structural properties and the categorization of visual inputs as bodies and body parts (Thierry *et al*., 2006; Urgesi *et al*., 2007). However, recent studies have reported a modulation of the N190 by the socio-emotional content of the stimuli (Borhani *et al*., 2015). Furthermore, in the face-processing literature, the N170 component (arguably reflecting similar visual processing as the N190) has been shown to be process-sensitive rather than purely category-sensitive (Blau *et al*., 2007; Hinojosa *et al*., 2015). Several studies report indeed a larger N170 for fearful, angry, and happy faces (see Hinojosa *et al*., 2015 and Schindler *et al*., 2020 for reviews). Our null result on the modulation of the N190 by pictures of hands conveying social or non-social affordances is therefore not purely in-line with the previous literature, but does not rule out early visual processing of sociality in hands. Indeed, our multivariate analyses show above-chance decoding of Shaking hands compared to Still ones from 132 ms on (see details below).

Crucially, we show that the Early Posterior Negativity (EPN) is differently influenced by the three hand-posture categories, with Shaking hands associated with the largest negative amplitude compared to the other two classes of stimuli. Previous EEG studies suggested that the EPN component indicates early perceptual tagging and prioritized visual processing (Olofsson *et al*., 2008), hence indexing natural selective attention for arousing and salient stimuli. Others highlighted how this component is strongly affected by negatively affective picture viewing (i.e., distressful scenes, threatening faces, gestures with negative valence, Junghöfer *et al*., 2001; Schupp, Öhman, *et al*., 2004; Schupp, Junghöfer, *et al*., 2004; Flaisch *et al*., 2009; Flaisch *et al*., 2011), compared to neutral images. However, when comparing EPN amplitudes during the processing of pleasant (e.g., erotic) and unpleasant (e.g., mutilation) stimuli with similar degrees of arousal, a bias in EPN modulation towards pleasant stimuli has been observed (Farkas and Sabatinelli, 2019). These findings have been framed within the perspective of understanding emotions as action sets (Lang *et al*., 2013), such that, for example, negative visual cues (i.e., insulting gestures, negative facial expressions) or pleasant and arousing scenes elicit preparation for immediate action. Coherently, some studies identified the left superior parietal cortex as the EPN main generator (Jaspers-Fayer *et al*., 2012). This area is also known to mediate visuo-motor transformations (Iacoboni, 2006; Mutha *et al*., 2011) and is fundamental to mediate interpersonal motor coordination (Sacheli *et al*., 2015; Era *et al*., 2018; Era *et al*., 2020). Nevertheless, other studies have identified the source of the EPN in extrastriate areas (Schupp *et al*., 2006, Frühholz *et al*., 2011). Here, by showing that the amplitude of the EPN component is sensitive to social content without negative valence, we reinforce the idea that the capture of visual attention is related to a more general form of preparation that drives attentional states, such as socially salient gestures.

### Occipito-parietal electrodes contribute to the decoding of hand gestures conveying social affordances

The univariate results on the modulation of the N190 and EPN by the social affordance conveyed by hand postures are complemented by our multivariate analyses. Our binary temporal decoding results confirm that salient information conveyed by hands is encoded in the brain at different stages. First, the classifier ‘s performance showed that when classifying Still vs Shaking hands (classification across time - Figure 3A), decoding accuracy was higher than chance at an early stage, starting earlier than 200 ms following stimulus presentation (as early as 132 ms after stimulus onset), at a later mid-latency stage in the 250-350 ms range and, finally, at a later stage up to 500 ms. This suggests that the neural patterns related to the processing of different hand stimuli conveying different levels of motion and social information are distinguishable early in the brain, a piece of information that was not clearly observable from the univariate analysis, demonstrating a slightly different sensitivity between univariate and multivariate approaches (as suggested by Grootswagers *et al*., 2017). The analysis of individuals ‘ peak decoding data also revealed larger accuracy values for Still vs Shaking decoding compared to accuracy values for Still vs Grasping, suggesting that the processing of socially relevant stimuli results in specific EEG patterns. Furthermore, our temporal generalization analysis showed that the patterns occurring in the first two time-windows are not the same, since the classifier ‘s performance did not generalise across time points, whereas the generators activated during the first time-window are likely recalled from 350 ms on (Figure 3B). Altogether, the binary temporal decoding approach complements the ERP analysis by adding evidence for different phenomena associated with the visual presentation of hands and suggests early process selectivity during the encoding of hands.

Moreover, the searchlight analysis shows a major contribution of occipito-temporal sites to the classification (Figure 4B and 4C), which are known to encode social information from bodily visual stimuli (Lingnau and Downing, 2015). The classifier ‘s performance resulted in significantly above-chance level at around 150 ms also when classifying Grasping vs Shaking (classification across time and electrodes, Figure 4C). These results suggest that early processing (<200 ms), usually linked to the sole categorization of visual stimuli appears to be sensitive to both implied motion and social information. Furthermore, while the analysis of early ERP amplitudes (Figure 2) and peak decoding latency of our time decoding did not reach significance (Figure 3D), the cluster-based analysis on the topographical distribution of classification weights from our Classification across time and electrodes shows that the presence of social information in hands is associated with significant early categorization over occipito-temporal and fronto-central sites (Figure 4D and 4E). Therefore, future studies aiming at revealing early modulation of visual activity by social content in bodily stimuli should focus both on the ERPs associated with the perception of social information and on MVPA decoding approaches.

In the case of hands, previous work focusing on the LOTC during hand action discrimination with and without social meaning showed that patterns in this hub allowed the discrimination of hand postures (Bracci *et al*., 2018). However, the authors ‘ conclusions pointed towards a category-selective (i.e., capable of discriminating body parts) role of the LOTC. Here, our results suggest a potential process-selective phenomenon (i.e., capable of representing the meaning of actions) over occipito-temporal sites. Indeed, our analyses distinguished between hands with and without implied actions (i.e., Still vs Grasping/Shaking), and crucially, Shaking hands observation shows distinct neural patterns compared to Grasping images, therefore suggesting that social actions might be encoded differently compared to actions towards objects.

Finally, higher-level visual areas are also integrated into a wider fronto-parietal network involved in action understanding and execution, where the activation of frontal and motor areas is thought to facilitate visual perception via top-down connections (Kilner and Frith, 2007; Bressler *et al*., 2008). Here, as revealed by the classification over time and electrodes (Figure 4A-C), the decoding of Shaking hands against Still hands shows that, in addition to the occipito-temporal sites, the performance of the classifier strongly relied on frontal and parietal electrodes. As EEG is known to have a poor spatial resolution, direct interpretation of the localization of these clusters of electrodes is limited. However, this could suggest top-down modulation processes from motor and parietal areas to the visual processing of hand postures conveying salient, social affordances compared to Still ones. On the other hand, only occipito-parietal sites significantly contributed to the classifier ‘s performance when differentiating Shaking hands from Grasping hands (i.e., both with implied motion). These results are coherent with previous studies on the role of the parietal cortex in understanding the goal of observed actions in primates (Fogassi *et al*., 2005; Filippini *et al*., 2018) and humans (Rizzolatti and Craighero, 2004; Hamilton and Grafton, 2006) and with the suggestion that others ‘ actions may represent social affordances, implying the selection of potential motor responses in parietal cortices (Orban *et al*., 2021). Furthermore, using fMRI-multivoxel pattern analysis, Wurm and colleagues (2017) have previously reported that the sociality of hand actions could be accurately decoded within the lateral occipital cortex (Wurm *et al*., 2017). Our results here complement this finding, by showing that the decoding of social and non-social implied actions by hands relies on occipito-parietal and occipito-temporal electrodes and provides new information regarding the precise timings of this process.

## Limitations

The interpretation of the results is limited by the use of static images. Although this choice was made to ensure EEG data quality, recent evidence showed that the use of dynamic stimuli only has minimal impact on signal-to-noise ratio and even reduces eye-artifact (Welke and Vessel, 2022). Future studies will aim at validating these results with videos to increase our understanding of how the brain categorizes social information.

## Conclusion

To sum up, we combined univariate and multivariate analyses to tackle the different dimensions of the EEG activity associated with the visual processing of hand gestures implying social affordances. This approach allowed us to integrate multiple sources of information, which together provide new evidence on the sensitivity of the EPN component in discriminating sociality in hand gestures and suggest that the integration of motion and social information happens in the early stages of visual processing, as suggested by our decoding analyses.

## SUPPLEMENTARY MATERIALS

### ERP analysis per each volunteer ‘s identity

Following one reviewer ‘s suggestion, we went back to our raw data wanting to evaluate if stimuli with spread-out fingers would show a different pattern than others. Therefore, we plotted and analyzed the ERPs separately for each identity of the volunteers whose hands were part of our stimuli set. Crucially, we saw that one particular stimulus, with a different position of the fingers for its Shaking picture, resulted in a strong reduction of the N190 compared to all other stimuli and all other identities (see Figure S1 - C). Since this is a potential confound that could influence the effects on the N190, as well as our trial-by-trial multivariate analysis, we have removed data linked to this person ‘s stimuli.

**Figure S1.**
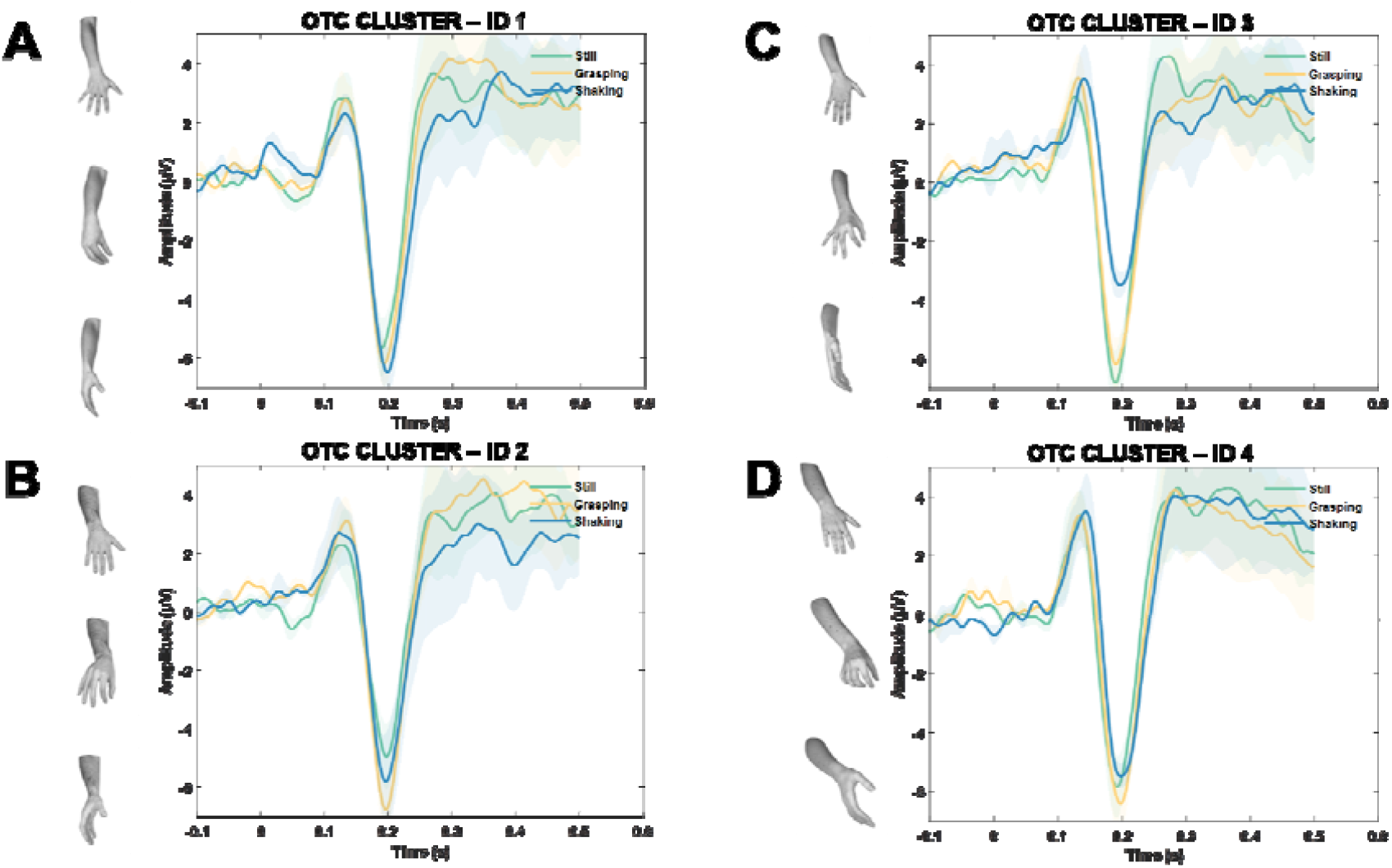
N190 and Early Posterior Negativity (EPN) event-related potentials over the bilateral occipito-temporal cluster (P7, PO7, P8, and PO8) for each of the four identities of the volunteers participating in our stimuli creation. We removed from all our main analyses the trials where stimuli belonging to identity 3 (Panel C) were shown.

## References

Argyle, M. (1972). Non-verbal communication in human social interaction. In: Non-verbal communication. Oxford, England: Cambridge U. Press, p. xiii, 443–xiii, 443.

Astafiev, S.V., Stanley, C.M., Shulman, G.L., et al. (2004). Extrastriate body area in human occipital cortex responds to the performance of motor actions. Nature Neuroscience, 7, 542–48

Avenanti, A., Candidi, M., Urgesi, C. (2013). Vicarious motor activation during action perception: beyond correlational evidence. Frontiers in Human Neuroscience, 7

Blau, V.C., Maurer, U., Tottenham, N., McCandliss, B.D. (2007). The face-specific N170 component is modulated by emotional facial expression. Behavioral and Brain Functions, 3.

Borhani, K., Làdavas, E., Maier, M.E., et al. (2015). Emotional and movement-related body postures modulate visual processing. Social Cognitive and Affective Neuroscience, 10, 1092–1101

Bracci, S., Cavina-Pratesi, C., Ietswaart, M., et al. (2012). Closely overlapping responses to tools and hands in left lateral occipitotemporal cortex. Journal of Neurophysiology, 107, 1443–56

Bracci, S., Ietswaart, M., Peelen, M.V., et al. (2010). Dissociable Neural Responses to Hands and Non-Hand Body Parts in Human Left Extrastriate Visual Cortex. Journal of Neurophysiology, 103, 3389–97

Bracci, S., Caramazza, A., Peelen, M.V. (2018). View-invariant representation of hand postures in the human lateral occipitotemporal cortex. NeuroImage, 181, 446–452

Bressler, S.L., Tang, W., Sylvester, C.M., et al. (2008). Top-Down Control of Human Visual Cortex by Frontal and Parietal Cortex in Anticipatory Visual Spatial Attention. The Journal of Neuroscience, 28, 10056–61

Campbell, J.I.D., Thompson, V.A. (2012). MorePower 6.0 for ANOVA with relational confidence intervals and Bayesian analysis. Behavior Research Methods, 44, 1255–65

Candidi, M., Stienen, B.M.C., Aglioti, S.M., et al. (2011). Event-Related Repetitive Transcranial Magnetic Stimulation of Posterior Superior Temporal Sulcus Improves the Detection of Threatening Postural Changes in Human Bodies. The Journal of Neuroscience, 31, 17547–54

Candidi, M., Stienen, B.M.C., Aglioti, S.M., et al. (2015). Virtual lesion of right posterior superior temporal sulcus modulates conscious visual perception of fearful expressions in faces and bodies. Cortex; a Journal Devoted to the Study of the Nervous System and Behavior, 65, 184–94

Candidi, M., Urgesi, C., Ionta, S., et al. (2008). Virtual lesion of ventral premotor cortex impairs visual perception of biomechanically possible but not impossible actions. Social Neuroscience, 3, 388–400

Cattaneo, L., Sandrini, M., Schwarzbach, J. (2010). State-dependent TMS reveals a hierarchical representation of observed acts in the temporal, parietal, and premotor cortices. Cerebral Cortex (New York, N.Y.: 1991), 20, 2252–58

Delorme, A., Makeig, S. (2004). EEGLAB: an open source toolbox for analysis of single-trial EEG dynamics including independent component analysis. Journal of Neuroscience Methods, 134, 9–21

Downing, P.E., Jiang, Y., Shuman, M., et al. (2001). A cortical area selective for visual processing of the human body. Science (New York, N.Y.), 293, 2470–73

van Driel, J., Olivers, C.N.L., Fahrenfort, J.J. (2021). High-pass filtering artifacts in multivariate classification of neural time series data. Journal of Neuroscience Methods, 352, 109080

Era, V., Aglioti, S.M., Candidi, M. (2020). Inhibitory Theta Burst Stimulation Highlights the Role of Left aIPS and Right TPJ during Complementary and Imitative Human–Avatar Interactions in Cooperative and Competitive Scenarios. Cerebral Cortex, 30, 1677–87

Era, V., Candidi, M., Gandolfo, M., et al. (2018). Inhibition of left anterior intraparietal sulcus shows that mutual adjustment marks dyadic joint-actions in humans. Social Cognitive and Affective Neuroscience, 13, 492–500

Espírito Santo, M.G., Maxim, O.S., Schürmann, M. (2017). N1 responses to images of hands in occipito-temporal event-related potentials. Neuropsychologia, 106, 83–89

Fan, R., Chang, K., Hsieh, C., et al. (2008). LIBLINEAR: A Library for Large Linear Classification

Farkas, A.H., Oliver, K.I., Sabatinelli, D. (2019). Emotional and feature-based modulation of the early posterior negativity. Psychophysiology, 57

Filippini, M., Breveglieri, R., Hadjidimitrakis, K., et al. (2018). Prediction of Reach Goals in Depth and Direction from the Parietal Cortex. Cell Reports, 23, 725–32

Flaisch, T., Häcker, F., Renner, B., et al. (2011). Emotion and the processing of symbolic gestures: an event-related brain potential study. Social Cognitive and Affective Neuroscience, 6, 109–18

Flaisch, T., Schupp, H.T. (2013). Tracing the time course of emotion perception: the impact of stimulus physics and semantics on gesture processing. Soc Cogn Affect Neurosci. 8, 820–7.

Flaisch, T., Schupp, H.T., Renner, B., et al. (2009). Neural systems of visual attention responding to emotional gestures. NeuroImage, 45, 1339–46

Fogassi, L., Ferrari, P.F., Gesierich, B., et al. (2005). Parietal Lobe: From Action Organization to Intention Understanding. Science, 308, 662–67

Frühholz, S., Jellinghaus, A., Herrmann, M. (2011). Time course of implicit processing and explicit processing of emotional faces and emotional words. Biol Psychol. 87, 265–74.

Gandolfo, M., Downing, P.E. (2019). Causal Evidence for Expression of Perceptual Expectations in Category-Selective Extrastriate Regions. Current Biology, 29, 2496–2500.e3

Goodale, M.A. (2014). How (and why) the visual control of action differs from visual perception. Proceedings of the Royal Society B: Biological Sciences, 281, 20140337

Grootswagers, T., Wardle, S.G., Carlson, T.A. (2017). Decoding Dynamic Brain Patterns from Evoked Responses: A Tutorial on Multivariate Pattern Analysis Applied to Time Series Neuroimaging Data. Journal of Cognitive Neuroscience, 29, 677–97

Gschwind, M., Pourtois, G., Schwartz, S., et al. (2012). White-Matter Connectivity between Face-Responsive Regions in the Human Brain. Cerebral Cortex, 22, 1564–76

Hamilton, A.F. de C., Grafton, S.T. (2006). Goal Representation in Human Anterior Intraparietal Sulcus. Journal of Neuroscience, 26, 1133–37

Haxby, J.V., Hoffman, E.A., Gobbini, M.I. (2002). Human neural systems for face recognition and social communication. Biological Psychiatry, 51, 59–67

Haxby, J.V., Hoffman, E.A., Gobbini, M.I. (2000). The distributed human neural system for face perception. Trends in Cognitive Sciences, 4, 223–33

Hinojosa, J.A., Mercado, F., Carretié, L. (2015). N170 sensitivity to facial expression: a meta-analysis. Neuroscience and Biobehavioral Reviews, 55, 498–509

Hugenberg, K., Wilson, J.P. (2013). Faces are central to social cognition. In: The Oxford handbook of social cognition. Oxford library of psychology. New York, NY, US: Oxford University Press, p. 167–93.

Iacoboni, M. (2006). Visuo-motor integration and control in the human posterior parietal cortex: evidence from TMS and fMRI. Neuropsychologia, 44, 2691–99

Ishizu, T., Amemiya, K., Yumoto, M., et al. (2010). Magnetoencephalographic study of the neural responses in body perception. Neuroscience Letters, 481, 36–40

Jaspers-Fayer, F., Ertl, M., Leicht, G., et al. (2012). Single-trial EEG–fMRI coupling of the emotional auditory early posterior negativity. NeuroImage, 62, 1807–14

Jung, T.P., Makeig, S., Humphries, C., et al. (2000). Removing electroencephalographic artifacts by blind source separation. Psychophysiology, 37, 163–78

Junghöfer, M., Bradley, M.M., Elbert, T.R., et al. (2001). Fleeting images: A new look at early emotion discrimination. Psychophysiology, 38, 175–78

Keselman, H.J., Rogan, J.C., Mendoza, J.L., et al. (1980). Testing the validity conditions of repeated measures F tests. Psychological Bulletin, 87, 479–81

King, J. R., &Dehaene, S. (2014). Characterizing the dynamics of mental representations: the temporal generalization method. Trends in cognitive sciences, 18(4), 203–210.

Kilner, J.M., Frith, C.D. (2007). A possible role for primary motor cortex during action observation. Proceedings of the National Academy of Sciences, 104, 8683–84

Lang, P.J., Simons, R.F., Balaban, M., et al. (2013). Attention and Orienting: Sensory and Motivational Processes. Psychology Press.

Lawrence, M. ez: Easy analysis and visualization of factorial experiments [online]. Available from: https://scholar.google.com/citations?view_op=view_citation&hl=en&user=7gTDUWMAAAAJ&citation_for_view=7gTDUWMAAAAJ:WF5omc3nYNoC [Accessed July 15, 2021].

Lingnau, A., Downing, P.E. (2015). The lateral occipitotemporal cortex in action. Trends in Cognitive Sciences, 19, 268–77

Mares, I., Ewing, L., Farran, E. K., Smith, F. W., & Smith, M. L. (2020). Developmental changes in the processing of faces as revealed by EEG decoding. Neuroimage, 211, 116660.

Maris, E., Oostenveld, R. (2007). Nonparametric statistical testing of EEG- and MEG-data. Journal of Neuroscience Methods, 164, 177–90

Meeren, H.K.M., Gelder, B. de, Ahlfors, S.P., et al. (2013). Different Cortical Dynamics in Face and Body Perception: An MEG study. PLOS ONE, 8, e71408

Milner, A.D., Goodale, M.A. (2008). Two visual systems re-viewed. Neuropsychologia, 46, 774–85

Milner, D., Goodale, M. (2006). The Visual Brain in Action. OUP Oxford.

Moreau, Q., Parrotta, E., Era, V., et al. (2019). Role of the occipito-temporal theta rhythm in hand visual identification. Journal of Neurophysiology, 123, 167–77

Moreau, Q., Pavone, E.F., Aglioti, S.M., et al. (2018). Theta synchronization over occipito-temporal cortices during visual perception of body parts. European Journal of Neuroscience, 48, 2826–35

Moro, V., Pernigo, S., Avesani, R., et al. (2012). Visual body recognition in a prosopagnosic patient. Neuropsychologia, 50, 104–17

Moro, V., Urgesi, C., Pernigo, S., et al. (2008). The Neural Basis of Body Form and Body Action Agnosia. Neuron, 60, 235–46

Mutha, P.K., Sainburg, R.L., Haaland, K.Y. (2011). Left Parietal Regions Are Critical for Adaptive Visuomotor Control. The Journal of Neuroscience, 31, 6972–81

Olofsson, J.K., Nordin, S., Sequeira, H., et al. (2008). Affective picture processing: An integrative review of ERP findings. Biological Psychology, 77, 247–65

Oostenveld, R., Fries, P., Maris, E., et al. (2011). FieldTrip: Open source software for advanced analysis of MEG, EEG, and invasive electrophysiological data. Computational Intelligence and Neuroscience, 2011, 156869

Orban, G.A., Lanzilotto, M., Bonini, L. (2021). From Observed Action Identity to Social Affordances. Trends in Cognitive Sciences, 25, 493–505

Orlov, T., Makin, T.R., Zohary, E. (2010). Topographic Representation of the Human Body in the Occipitotemporal Cortex. Neuron, 68, 586–600

Papeo, L., 2020. Twos in human visual perception. Cortex 132, 473–478.

Pitcher, D., Ungerleider, L.G. (2021). Evidence for a Third Visual Pathway Specialized for Social Perception. Trends in Cognitive Sciences, 25, 100–110

Pourtois, G., Peelen, M.V., Spinelli, L., Seeck, M., Vuilleumier, P. (2007). Direct intracranial recording of body-selective responses in human extrastriate visual cortex. Neuropsychologia, 45, 2621–2625

Ramsey, R. (2018). Neural Integration in Body Perception. Journal of Cognitive Neuroscience, 30, 1442–51

Rizzolatti, G., Craighero, L. (2004). The mirror-neuron system. Annual Review of Neuroscience, 27, 169–92

Sacheli, L.M., Candidi, M., Era, V., et al. (2015). Causative role of left aIPS in coding shared goals during human–avatar complementary joint actions. Nature Communications, 6, 7544

Sadeh, B., Pitcher, D., Brandman, T., et al. (2011). Stimulation of category-selective brain areas modulates ERP to their preferred categories. Current biology: CB, 21, 1894–99

Sato, W., Kochiyama, T., Yoshikawa, S., et al. (2001). Emotional expression boosts early visual processing of the face: ERP recording and its decomposition by independent component analysis. Neuroreport, 12, 709–14

Schindler, S., Bublatzky, F. (2020). Attention and emotion: an integrative review of emotional face processing as a function of attention. Cortex, 130, 362–386

Schupp, H.T., Junghöfer, M., Weike, A.I., et al. (2004). The selective processing of briefly presented affective pictures: An ERP analysis. Psychophysiology, 41, 441–49

Schupp, H.T., Öhman, A., Junghöfer, M., et al. (2004). The Facilitated Processing of Threatening Faces: An ERP Analysis. Emotion, 4, 189–200

Schupp, H.T., Flaisch, T., Stockburger, J., Junghofer, M. (2006). Emotion and attention: event-related brain potential studies. Progress in Brain Research, 156, 31–51

Schupp, H.T., Stockburger, J., Bublatzky, F., Junghofer, M., Weike, A.I., Hamm, A.O. (2007). Explicit attention interferes with selective emotion processing in human extrastriate cortex. BMC Neuroscience, 8

Sliwinska, M.W., Bearpark, C., Corkhill, J., et al. (2020). Dissociable pathways for moving and static face perception begin in early visual cortex: Evidence from an acquired prosopagnosic. Cortex; a Journal Devoted to the Study of the Nervous System and Behavior, 130, 327–39

Smith, F.W., Smith, M.L. (2019). Decoding the dynamic representation of facial expressions of emotion in explicit and incidental tasks. NeuroImage, 195, 261–71

Stewart, A.X., Nuthmann, A., Sanguinetti, G. (2014). Single-trial classification of EEG in a visual object task using ICA and machine learning. Journal of Neuroscience Methods, 228, 1–14

Tarhan, L., Konkle, T. (2020). Sociality and interaction envelope organize visual action representations. Nature Communications, 11, 3002

Taylor, J.C., Downing, P.E. (2011). Division of Labor between Lateral and Ventral Extrastriate Representations of Faces, Bodies, and Objects. Journal of Cognitive Neuroscience, 23, 4122–37

Taylor, J.C., Wiggett, A.J., Downing, P.E. (2007). Functional MRI Analysis of Body and Body Part Representations in the Extrastriate and Fusiform Body Areas. Journal of Neurophysiology, 98, 1626–33

Thierry, G., Pegna, A.J., Dodds, C., et al. (2006). An event-related potential component sensitive to images of the human body. NeuroImage, 32, 871–79

Treder, M.S. (2020). MVPA-Light: A Classification and Regression Toolbox for Multi-Dimensional Data. Frontiers in Neuroscience, 0

Tucciarelli, R., Turella, L., Oosterhof, N.N., et al. (2015). MEG Multivariate Analysis Reveals Early Abstract Action Representations in the Lateral Occipitotemporal Cortex. The Journal of Neuroscience: The Official Journal of the Society for Neuroscience, 35, 16034–45

Tucciarelli, R., Wurm, M., Baccolo, E., et al. (2019). The representational space of observed actions M. Lescroart, R. B. Ivry, M. Lescroart, et al. (eds). eLife, 8, e47686

Urgesi, C., Berlucchi, G., Aglioti, S.M. (2004). Magnetic stimulation of extrastriate body area impairs visual processing of nonfacial body parts. Current biology: CB, 14, 2130– 34

Urgesi, C., Calvo-Merino, B., Haggard, P., et al. (2007). Transcranial Magnetic Stimulation Reveals Two Cortical Pathways for Visual Body Processing. Journal of Neuroscience, 27, 8023–30

Urgesi, C., Candidi, M., Avenanti, A. (2014). Neuroanatomical substrates of action perception and understanding: an anatomic likelihood estimation meta-analysis of lesion-symptom mapping studies in brain injured patients. Frontiers in Human Neuroscience, 8

Vahid, A., Mückschel, M., Stober, S., et al. (2020). Applying deep learning to singletrial EEG data provides evidence for complementary theories on action control. Communications Biology, 3, 1–11

Varoquaux, G., Raamana, P.R., Engemann, D.A., et al. (2017). Assessing and tuning brain decoders: Cross-validation, caveats, and guidelines. NeuroImage, 145, 166–79

Welke, D., Vessel, E.A. (2022). Naturalistic viewing conditions can increase task engagement and aesthetic preference but have only minimal impact on EEG quality. NeuroImage, 56, 119218

Wurm, M.F., Caramazza, A. (2019). Lateral occipitotemporal cortex encodes perceptual components of social actions rather than abstract representations of sociality. NeuroImage, 202, 116153

Wurm, M.F., Caramazza, A., Lingnau, A. (2017). Action Categories in Lateral Occipitotemporal Cortex Are Organized Along Sociality and Transitivity. The Journal of Neuroscience, 37, 562–75

